# An Autoencoder and Artificial Neural Network-based Method to Estimate Parity Status of Wild Mosquitoes from Near-infrared Spectra

**DOI:** 10.1101/2020.01.25.919878

**Authors:** Masabho P. Milali, Samson S. Kiware, Nicodem J. Govella, Fredros Okumu, Naveen Bansal, Serdar Bozdag, Jacques D. Charlwood, Marta Maia, Sheila B. Ogoma, Floyd E. Dowell, George F. Corliss, Maggy T. Sikulu-Lord, Richard J. Povinelli

## Abstract

**Background:** After mating, female mosquitoes need animal blood to develop their eggs. In the process of acquiring blood, they may acquire pathogens, which may cause different diseases to humans such as malaria, zika, dengue, and chikungunya. Therefore, knowing the parity status of mosquitoes is useful in control and evaluation of infectious diseases transmitted by mosquitoes, where parous mosquitoes are assumed to be potentially infectious. Ovary dissections, which currently are used to determine the parity status of mosquitoes, are very tedious and limited to very few experts. An alternative to ovary dissections is near-infrared spectroscopy (NIRS), which can estimate the age in days and the infectious state of laboratory and semi-field reared mosquitoes with accuracies between 80 and 99%. No study has tested the accuracy of NIRS for estimating the parity status of wild mosquitoes.

**Methods and results:** In this study, we train artificial neural network (ANN) models on NIR spectra to estimate the parity status of wild mosquitoes. We use four different datasets: *An. arabiensis* collected from Minepa, Tanzania (Minepa-ARA); *An. gambiae* collected from Muleba, Tanzania (Muleba-GA); *An. gambiae* collected from Burkina Faso (Burkina-GA); and *An.gambiae* from Muleba and Burkina Faso combined (Muleba-Burkina-GA). We train ANN models on datasets with spectra preprocessed according to previous protocols. We then use autoencoders to reduce the spectra feature dimensions from 1851 to 10 and re-train ANN models. Before the autoencoder was applied, ANN models estimated parity status of mosquitoes in Minepa-ARA, Muleba-GA, Burkina-GA and Muleba-Burkina-GA with out-of-sample accuracies of 81.9 ± 2.8% (N=927), 68.7 ± 4.8% (N=140), 80.3 ± 2.0% (N=158), and 75.7 ± 2.5% (N=298), respectively. With the autoencoder, ANN models tested on out-of-sample data achieved 97.1 ± 2.2%, (N=927), 89.8 ± 1.7% (N=140), 93.3 ± 1.2% (N=158), and 92.7 ± 1.8% (N=298) accuracies for Minepa-ARA, Muleba-GA, Burkina-GA, and Muleba-Burkina-GA, respectively.

**Conclusion:** These results show that a combination of an autoencoder and an ANN trained on NIR spectra to estimate parity status of wild mosquitoes yields models that can be used as an alternative tool to estimate parity status of wild mosquitoes, especially since NIRS is a high-throughput, reagent-free, and simple-to-use technique compared to ovary dissections.

## Introduction

Evaluation of existing malaria control interventions such as insecticide-treated nets (ITNs) and indoor residual spraying (IRS) relies upon, among other factors, the assessment of the changes occurring in the mosquito parity structure prior to and after implementation of an intervention [1–3]. The parity status of mosquitoes corresponds with their capability to transmit *Plasmodium* parasites with an assumption that parous mosquitoes are more highly capable than nulliparous mosquitoes, as they may have accessed parasite-infected blood. A shift in the parity structure towards a population with more nulliparous mosquitoes signifies a reduction in the risk of disease transmission [1, 4, 5], as the chances that mosquitoes carry malaria parasite declines [6].

The current standard technique for estimating the parity status of female mosquitoes involves dissection of their ovaries to separate mosquitoes into those that have previously laid eggs, known as the parous group (assumed to be old and potentially infectious), and those that do not have a gonotrophic history, known as the nulliparous group (assumed to be young and non-infectious) [7]. Another standard technique also based on the dissection of ovaries determines the number of times a female mosquito has laid eggs [8]. However, both techniques are laborious, time consuming, and require skilled technicians. These technical difficulties lead to analysis of small sample sizes that often fail to capture the heterogeneity of a mosquito population.

Near infrared spectroscopy (NIRS) technology, complimented by techniques from machine learning, has been demonstrated to be an alternative tool for predicting age, species, and infectious status of laboratory and semi-field raised mosquitoes [9–20]. NIRS is a rapid, non-invasive, reagent-free technique that requires minimal skills to operate, allowing hundreds of samples to be analyzed in a day. However, the accuracy of NIRS techniques for predicting the parity status of wild mosquitoes has not been tested. Moreover, recently, it has been reported that models trained on NIR spectra using an artificial neural network (ANN) estimate the age of laboratory-reared *An. arabiensis, An.gambiae, Aedes aegypti*, and *Aedes albopictus* with accuracies higher than models trained on NIR spectra using partial least squares (PLS) [14].

In this study, we train ANN models on NIR spectra preprocessed according to an existing protocol [12] to estimate the parity status of wild *An. gambiae* s.s and *An. arabiensis*. We then apply autoencoders to reduce the spectra feature space from 1851 to 10 and re-train ANN models. The ANN model achieved an average accuracy of 72% and 93% before and after applying the autoencoder, respectively. These results strongly suggest ANN models trained on autoencoded NIR spectra as an alternative tool to estimate the parity status of wild *An. gambiae* and *An. arabiensis*. High-throughput, non-invasive, reagent free, and simple to use NIRS analysis complements the limitations of ovary dissections.

## Materials and Methods

### Ethics approvals

Ethics approvals for collecting mosquitoes in Minepa-ARA, Burkina-GA and Muleba-GA datasets from residents’ homes were obtained from Ethics Review Boards of the Ifakara Health Institute (IHI-IRB/No. 17-2015), the Colorado State University (approval No. 09-1148H), and the Kilimanjaro Christian Medical College (Certificate No. 781), respectively.

### Data

We use data from wild *An. arabiensis* (Minepa-ARA) collected from Minepa, a village in southeastern Tanzania (already published in [21] and available for reuse), from wild *An. gambiae s.s* (Muleba-GA) collected from Muleba, northwestern Tanzania (mosquitoes published in [22], permission to reuse obtained from the senior author), and from wild *An. gambiae s.s* collected from Bougouriba and Diarkadou-gou villages in Burkina Faso (Burkina-GA) (published in [10]and publicly available for reuse).

Mosquitoes in the Minepa-ARA and Muleba-GA datasets were captured using CDC light traps placed inside residential homes. Mosquitoes that were morphologically identified as members of the *Anopheles gambiae* complex were further processed. Prior to scanning, wild mosquitoes collected in Minepa were killed by freezing at −20°C for 20 minutes and left to re-equilibrate to room temperature for 30 minutes. Wild mosquitoes collected in Muleba were killed using 75% ethanol, dissected according to the technique described by Detinova [23] to determine their parity status, and preserved in silica gel. Mosquitoes in Minepa-ARA were dissected after scanning. Following a previous published protocol to collect spectra [12], mosquitoes in both Minepa-ARA and Muleba-GA were scanned using a LabSpec 5000 near-infrared spectrometer with an integrated light source (ASD Inc., Longmont, CO). After spectra collection, mosquitoes in Minepa-ARA were dissected to score their parity status. Then polymerase chain reaction (PCR) was conducted on DNA extracted from mosquito legs (in both Minepa-ARA and Muleba-GA) to identify species type as previously described [24]. Each mosquito was labeled with a unique identifier code linking each NIR spectrum to parity dissection and PCR information.

Data from wild *An. gambiae s.s* from Burkina Faso were published in [10] and publicly available for reuse. These mosquitoes are referred to as independent test sets 2 and 3 (ITS 2 and ITS 3) in [10]. ITS 2 has 40 nulliparous and 40 parous mosquitoes, and ITS 3 has 40 nulliparous and 38 parous mosquitoes. In this study, we combine these two datasets into one dataset and refer it as Burkina-GA. Mosquitoes in Burkina-GA (N = 158) were collected in 2013 in Burkina Faso from Bougouriba and Diarkadou-gou villages using either indoor aspiration or a human baited tent trap, and their ovaries were dissected according to the Detinova method [23]. Mosquitoes were preserved in silica gel before their spectra were collected using a LabSpec4i spectrometer (ASD Inc., Boulder, CO, USA).

### Model training and testing

We trained models on four datasets, namely Minepa-ARA, Muleba-GA, Burkina-GA, and Muleba-Burkina-GA (Muleba-GA and Burkina-GA combined). Before training models, spectra in all four datasets were pre-processed according to the previously published protocol [12] and divided into two groups (nulliparous and parous). Spectra in the nulliparous and parous groups were labeled zero and one, respectively. The two groups were then merged, randomized, and divided into a training set (75%; N = 927 for Minepa-ARA, N = 140 for the Muleba-GA, N = 158 for Burkina-GA and N = 298 for Muleba-Burkina-GA) and a test set (the remaining 25% in each dataset). On each dataset, using ten Monte-Carlo cross validations [14, 25] and Levenberg-Marquardt optimization, a one hidden layer, ten-neuron feed-forward ANN model with logistic regression as a transfer function was trained and tested in MATLAB (Fig 1).

**Figure 1.**
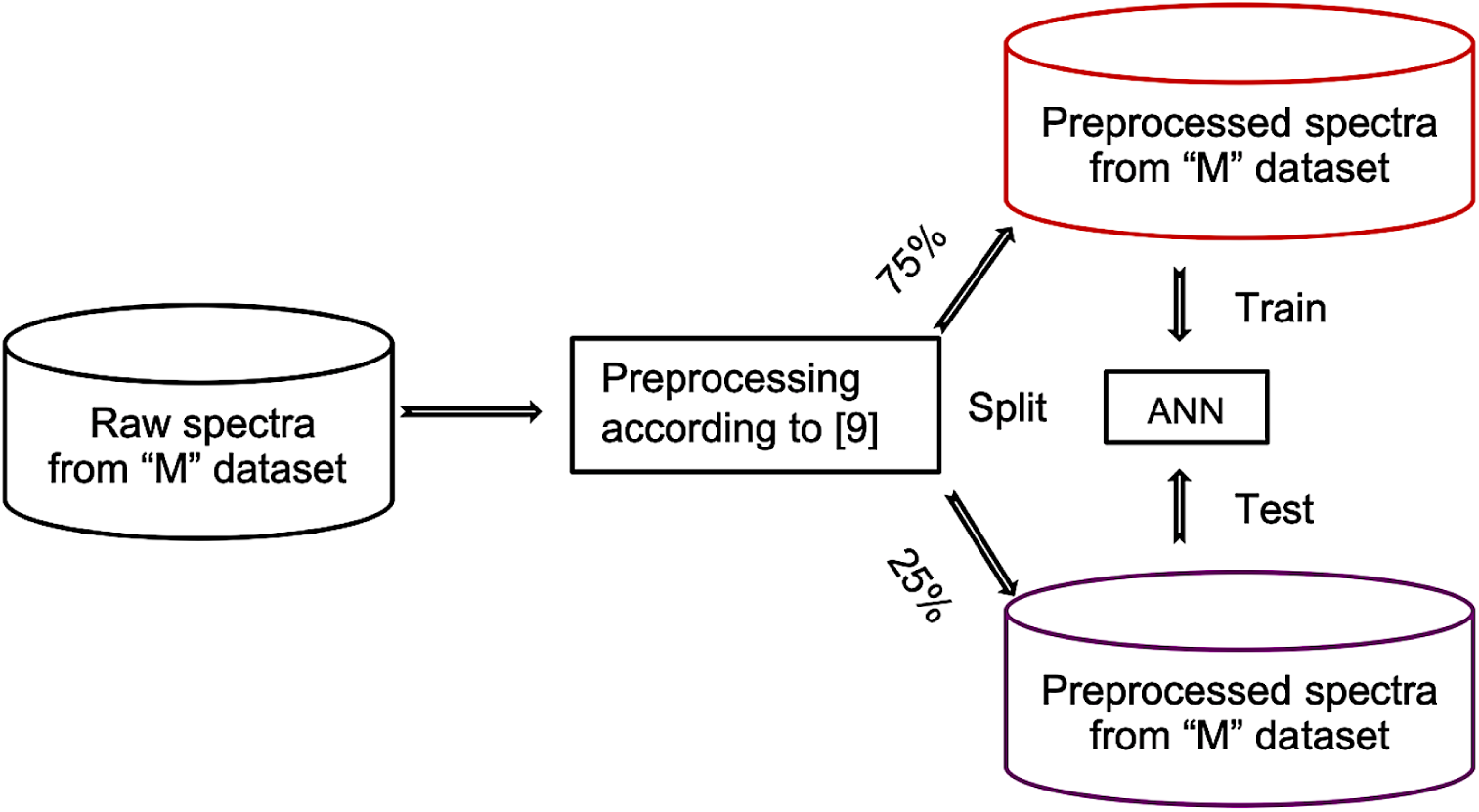
Training and testing ANN model on spectra preprocessed according to Mayagaya et al. [26]. ‘M’ is either Minepa-ARA, Muleba-GA, Burkina-GA, or Muleba-Burkina-GA

Based on the accuracy of the model presented in Tables 2 and 3, and Figs 7 and 8 in the Results and Discussion section, we explored how to improve the model accuracy. Normally a parous class, unlike a nulliparous class, often is represented by a limited number of samples, posing a problem of data imbalance during model training. In this case, a large amount of data is required to obtain enough samples in a parous class for a model to learn and characterize it accurately. Obtaining enough data for model training is always challenging. The most common ways of dealing with the data imbalance are either to discard samples from a nulliparous class to equal the number of samples in a parous class or to bootstrap samples in a parous class [27]. However, discarding data to equalize the data distribution in two classes in the training set leaves an imbalanced test set. Also, it is this imbalanced scenario to which the model will be applied in real cases. In addition, throwing away samples, especially from data sets with a high dimension feature space, can lead to over-fitting the model. Alternatively, for datasets with a high dimension feature space, instead of discarding data from a class with a large number of samples, feature reduction techniques are employed [27]. Feature reduction reduces the size of the hypothesis space initially presented in the original data, thereby reducing the size of data required to adequately train the model. Principal component analysis (PCA) and partial least squares (PLS) are the commonly used unsupervised and supervised feature reduction methods, respectively, especially for cases whose features are linearly related [28, 29]. Autoencoders recently are used as an alternative to PCA in cases involving both linear and non-linear relationships [30–33].

**Table 1.** Accuracies of reconstructing original feature spaces from encoded feature spaces. MSE = mean square error.

| Metric | Steps | Encoded-Minepa-ARA | Encoded-Muleba-GA | Encoded-Burkina-GA |
| --- | --- | --- | --- | --- |
| MSE | Step 1 | 0.0046 | 0.0029 | 0.0031 |
|  | Step 2 | 0.00005 | 0.0027 | 0.0022 |
|  | Step 3 | 0.00008 | 0.0029 | 0.0011 |

**Table 2.** Performance of an ANN model trained on 75% of mosquito spectra with 1851 features (before autoencoder) and tested on the remaining 25% spectra (out of the sample testing). Minepa-ARA (Nulliparous = 656, Parous = 271), Muleba-GA (Nulliparous = 119, Parous = 21) Burkina-GA (Nulliparous = 80, Parous = 78)

|  | Minepa-ARA<br>(N=927) | Muleba-GA<br>(N=140) | Burkina-GA<br>(N=158) | Muleba-Burkina-GA<br>(N=298) |
| --- | --- | --- | --- | --- |
| **Accuracy (%)** | $81.9 \pm 2.8$ | $68.7 \pm 4.8$ | $80.3 \pm 2.0$ | $75.7 \pm 2.5$ |
| **Sensitivity (%)** | $79.7 \pm 3.2$ | $37.8 \pm 6.6$ | $76.5 \pm 2.1$ | $70.2 \pm 3.1$ |
| **Specificity (%)** | $86.0 \pm 1.6$ | $80.1 \pm 2.7$ | $88.3 \pm 2.3$ | $77.6 \pm 2.9$ |
| **Precision (%)** | $74.3 \pm 3.4$ | $31.3 \pm 5.2$ | $77.8 \pm 1.8$ | $68.8 \pm 3.2$ |
| **AUC (%)** | 77.2 | 55.9 | 83.6 | 76.4 |

**Table 3.** Confusion matrices showing accuracies of the models in absolute values when the models were trained on spectra before feature reduction by autoencoder. A) Minepa-ARA, B) Muleba-GA, C) Burkina-GA and D) Muleba-Burkina-GA. Results from the last Monte-Carlo cross validation.

|  |  | Actual Parity |  |  |
| --- | --- | --- | --- | --- |
|  | Estimates | Nulliparous | Parous | Total |
| **A** | Nulliparous | 165 | 17 | 182 |
|  | Parous | 31 | 61 | 92 |
|  | Total | 196 | 78 | 274 |
| **B** | Nulliparous | 28 | 4 | 32 |
|  | Parous | 8 | 3 | 11 |
|  | Total | 36 | 7 | 43 |
| **C** | Nulliparous | 20 | 6 | 26 |
|  | Parous | 4 | 18 | 22 |
|  | Total | 24 | 24 | 48 |
| **D** | Nulliparous | 46 | 9 | 55 |
|  | Parous | 14 | 22 | 36 |
|  | Total | 60 | 31 | 91 |

**Figure 2.**
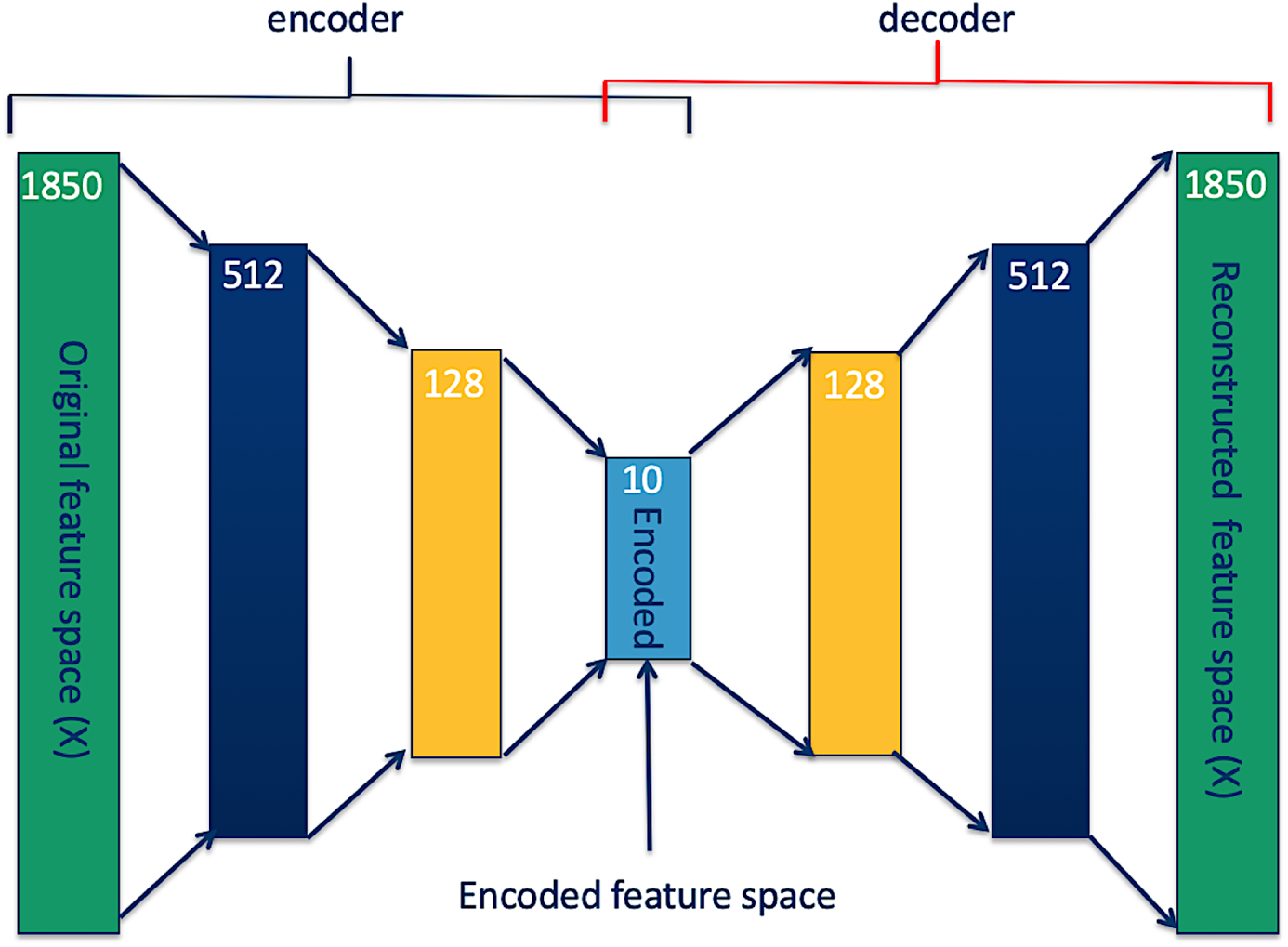
Autoencoders reducing feature space dimension.

**Figure 3.**
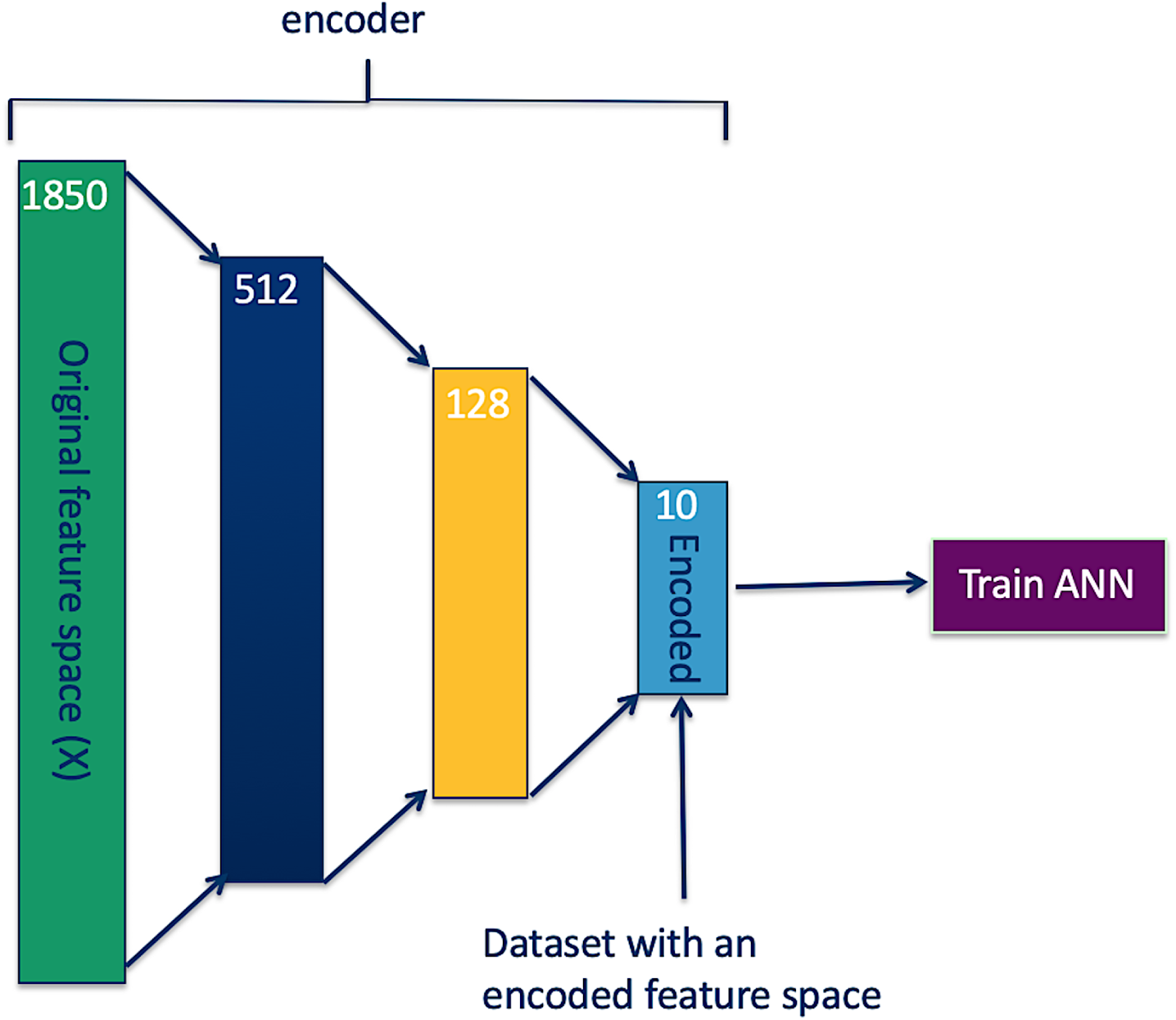
ANN model trained on a dataset with an encoded feature space.

**Figure 4.**
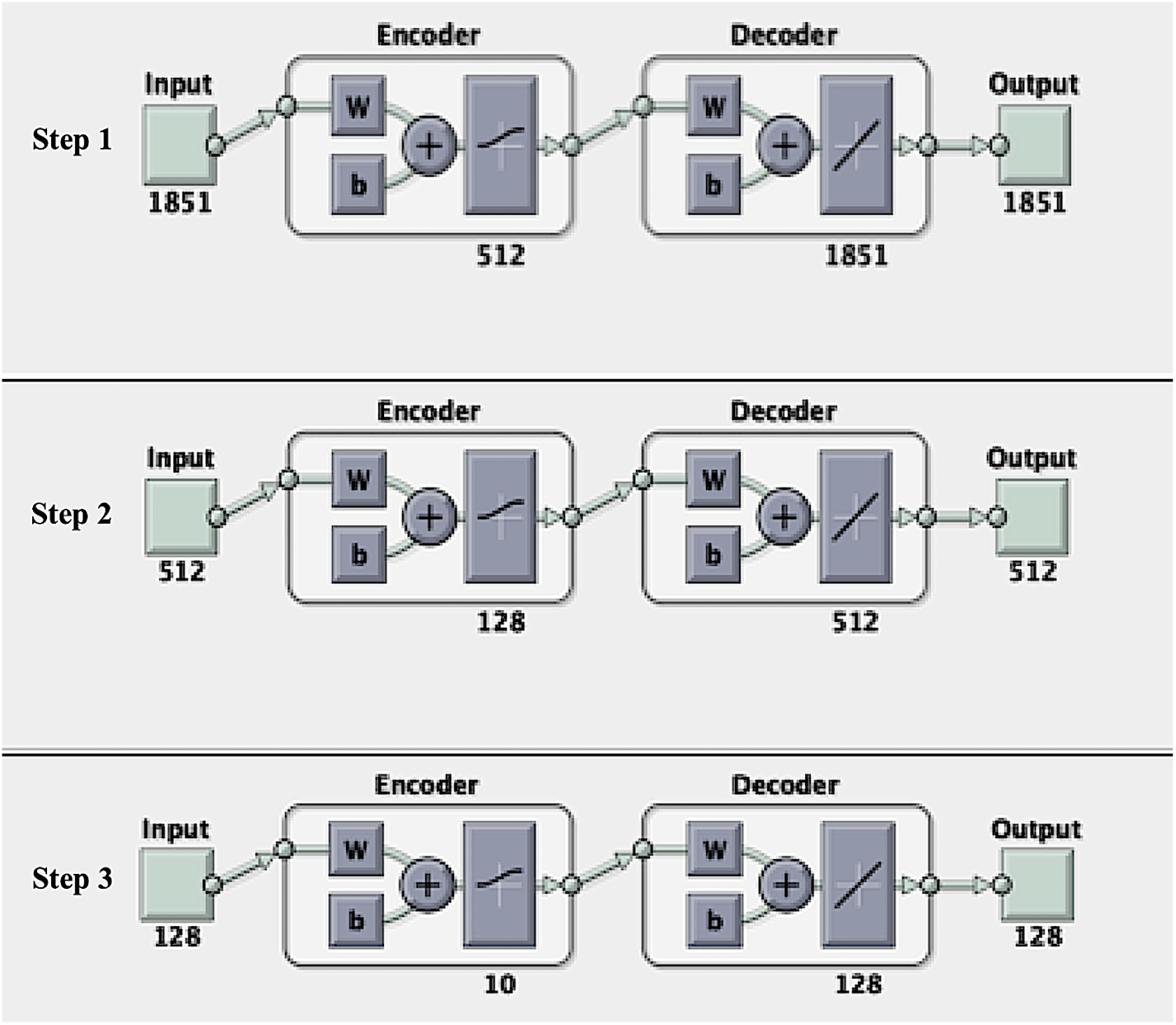
Reducing spectra feature space using an autoencoder and reconstructing original feature spaces from their respective encoded feature spaces (reconstruction accuracies presented in Table 1). Figures generated from MATLAB.

**Figure 5.**
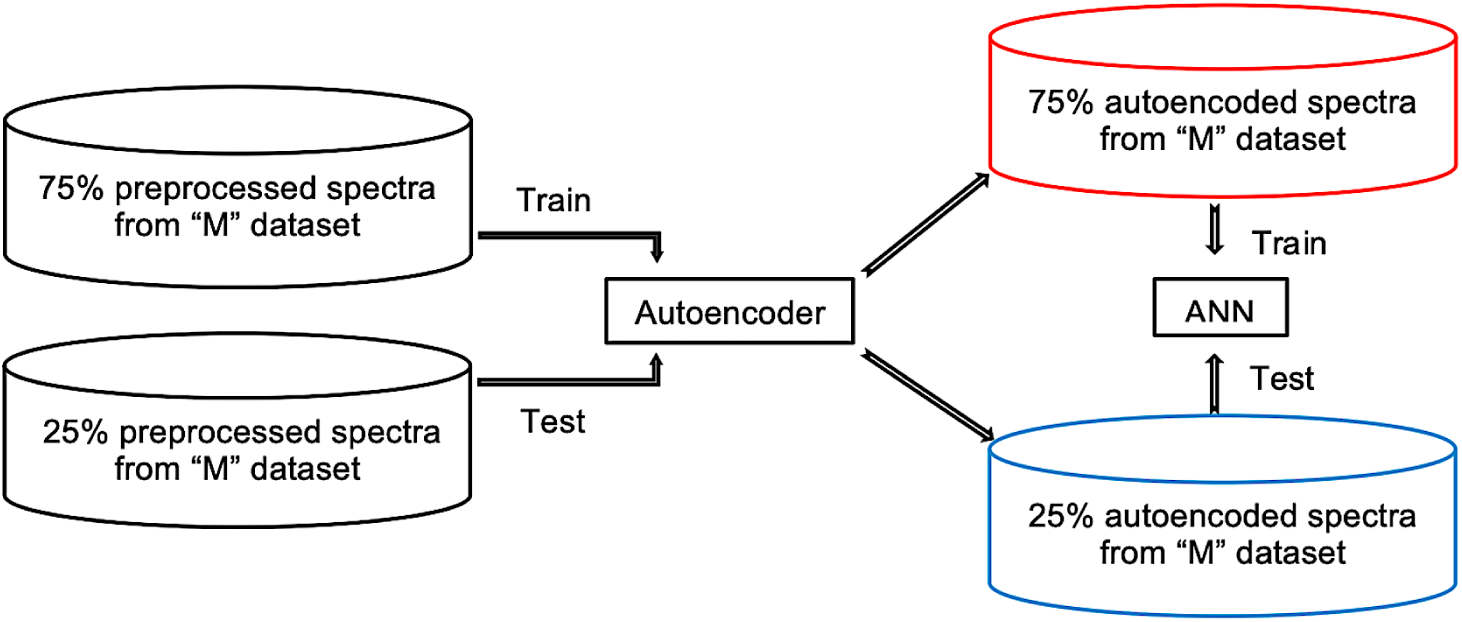
Training and testing of ANN model on autoencoded spectra. M is either Minepa-ARA, Muleba-GA, Burkina-GA, or Muleba-Burkina-GA dataset.

**Figure 6.**
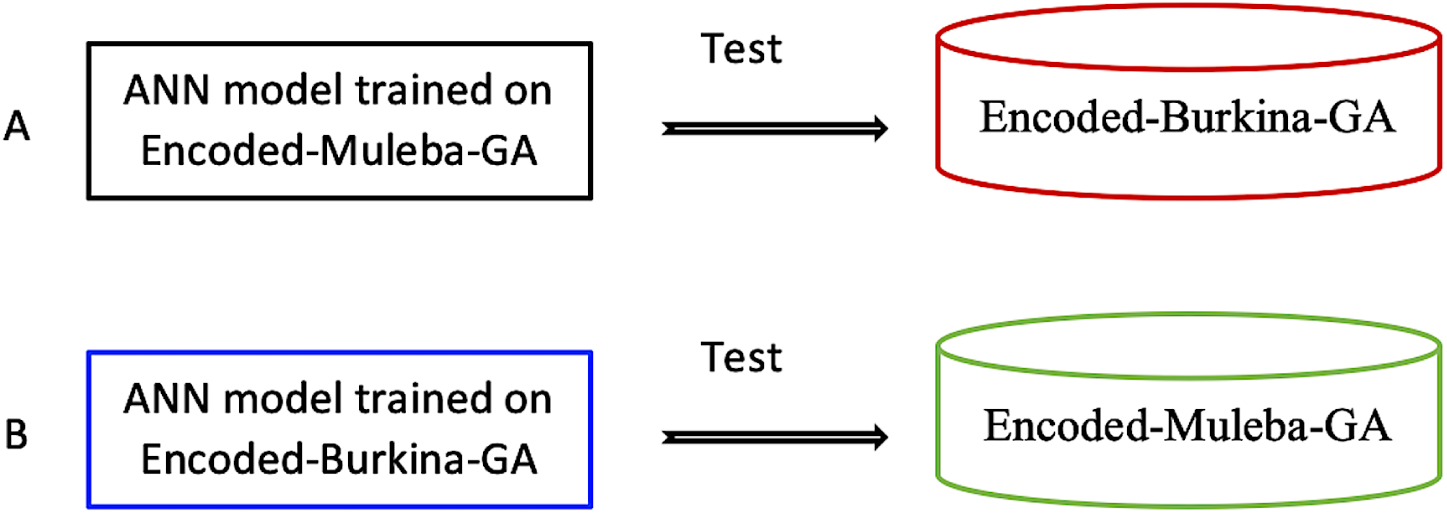
Independent testing of ANN model trained on encoded datasets. A) Applying ANN model trained on Encoded-Muleba-GA dataset to estimate the parity status of mosquitoes in autoencoded Burkina-GA dataset. B) Applying ANN model trained on Encoded-Burkina-GA dataset to estimate the parity status of mosquitoes in Encoded-Muleba-GA dataset

**Figure 7.**
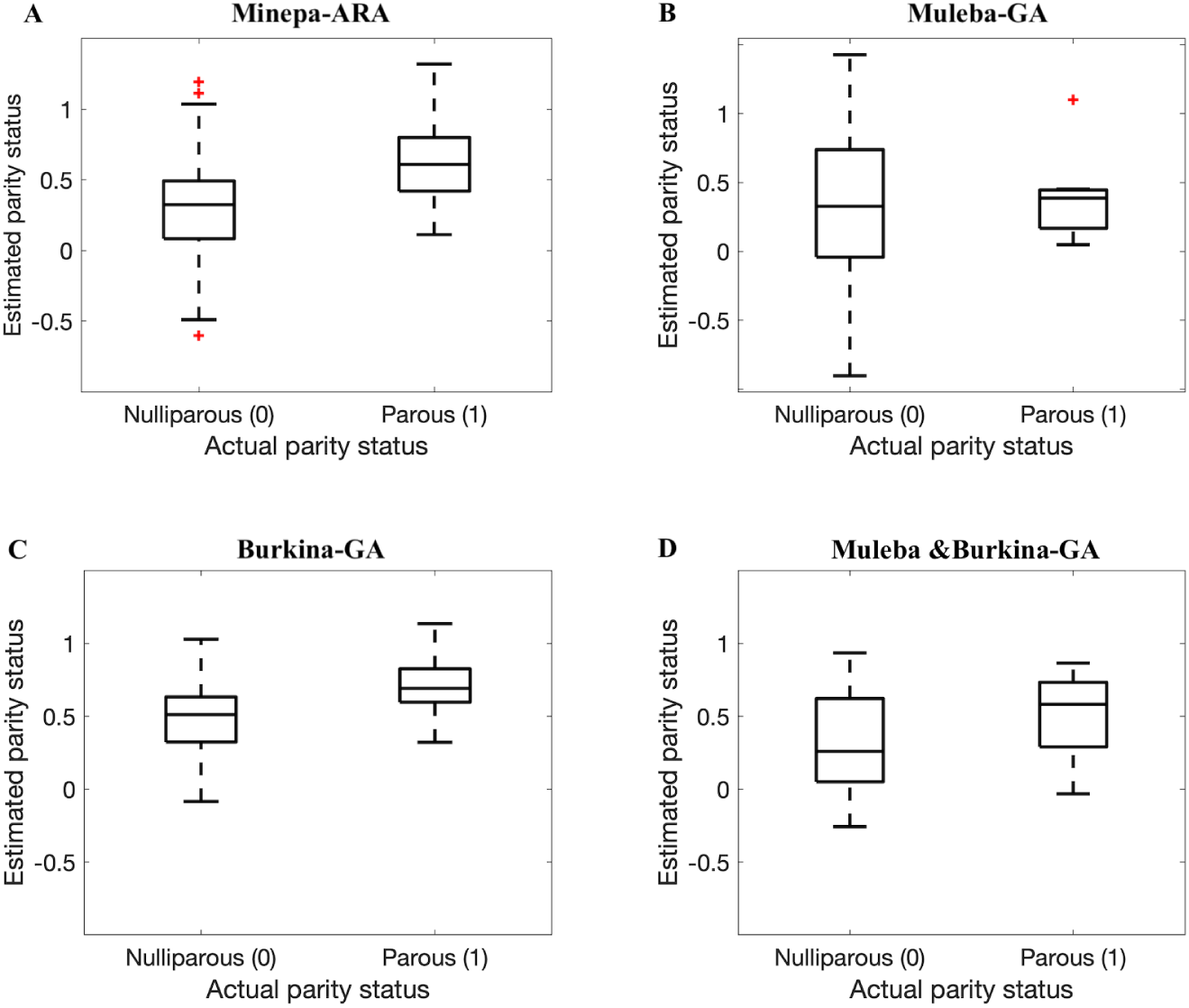
Box plots showing results when ANN models trained on 75% of spectra before the autoencoder was applied and tested on the remaining spectra (25%) (out of the sample testing). A, B, C, and D represent results for Minepa-ARA, Muleba-GA, Burkina-GA, and Muleba & Burkina-GA (mosquitoes in Muleba-GA and Burkina-GA datasets combined) datasets, respectively

**Figure 8.**
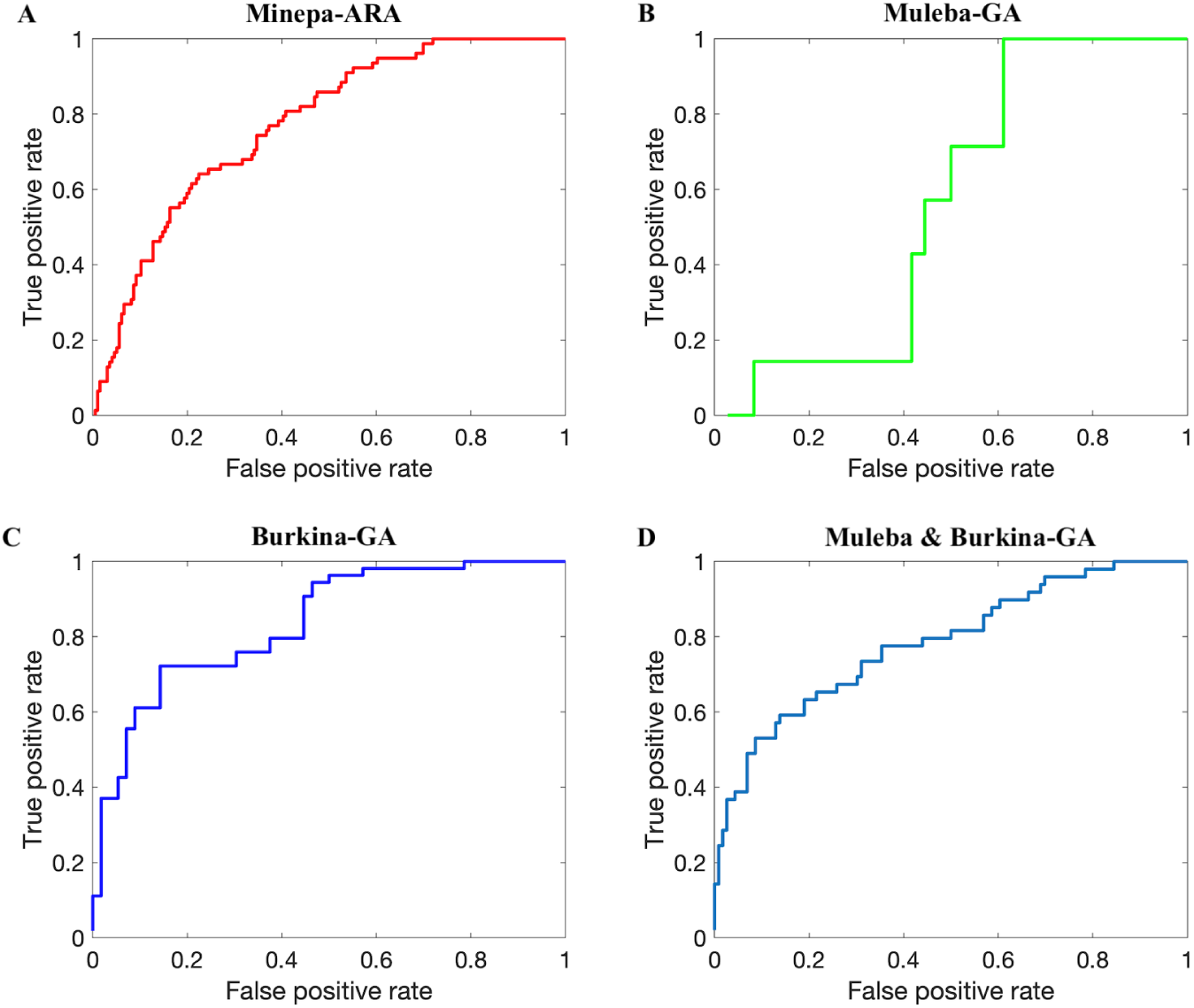
ROC curves (AUCs presented in the last row of Table 1) showing results when ANN models trained on 75% of spectra before the autoencoder was applied and tested on the remaining spectra (25%) (out of the sample testing). A, B, C, and D represent results for Minepa-ARA, Muleba-GA, Burkina-GA, and Muleba & Burkina-GA (mosquitoes in Muleba-GA and Burkina-GA datasets combined) datasets, respectively

An autoencoder is an unsupervised ANN that learns both linear and non-linear relationships present in data and represents them in a new reduced dimension data space (which also can be used to regenerate the original data space) without losing important information [34–36]. The autoencoder has two parts, the encoder part where an original dataset is encoded to a desired reduced feature space (encoded dataset) and the decoder part where the encoded dataset is decoded to an original dataset to determine how accurately the encoded dataset represents the original dataset. Fig 2 illustrates an example of an autoencoder in which an 1850-feature dataset is stepwise encoded to a 10-feature dataset. There is no formula for the number and size of steps to take to get to a desired feature size. However, taking several steps results on losing very little information, compared with taking a single step.

Once an encoded feature space can reconstruct the original feature space with an acceptable accuracy, the decoder is detached, and a desired model (in our case an ANN binary classifier) is trained on the encoded feature space as shown in Fig 3.

Egg laying appears to be affected by both linear and non-linear relationships. Hence, we separately train autoencoders on the Minepa-ARA, Muleba-GA, Burkina-GA, and Muleba-Burkina-GA datasets to reduce spectra feature dimensions from 1851 to 10 (Fig 4). Table 1 presents accuracies of reconstructing original feature spaces from their respective encoded feature spaces.

We refer to the autoencoded Minepa-ARA, Muleba-GA, Burkina-GA, and Muleba-Burkina-GA datasets as Encoded-Minepa-ARA, Encoded-Muleba-GA, Encoded-Burkina-GA, and Encoded-Muleba-Burkina-GA, respectively. We then train ANN models on Encoded-Minepa-ARA, Encoded-Muleba-GA, Encoded-Burkina-GA, and Encoded-Muleba-Burkina-GA (Fig 5).

Finally, we used the Encoded-Burkina-GA and the Encoded-Muleba-GA datasets as independent test sets to test accuracies of ANN models trained on the Encoded-Muleba-GA dataset and on the Encoded-Burkina-GA dataset, respectively (Figure 6A and B).

## Results and discussion

In this study, we demonstrated that near-infrared spectroscopy (NIRS) can estimate accurately the parity status of wild collected *An. arabiensis* and *An. gambiae* s.s. Referring to the published results in [14] (ANN models achieve higher accuracies than PLS models), we trained and tested an ANN model on NIRS spectra in four different datasets pre-processed according to a previous published protocol [12]. The model achieved accuracies between 55.9 and 81.9% (Tables 2 and 3, Figs 7 and 8). Table 2 further presents various metrics to score performance of our classifiers, namely sensitivity, specificity, precision, and area under the receiver operating characteristic (ROC) curve (AUC). We calculated sensitivity, specificity, accuracy, and precision of the model using Equations 1, 2, 3, and 4, respectively [37–40].

Let

- TP = Number of mosquitoes correctly classified by the model as parous,
- FP = Number of mosquitoes wrongly classified by the model as parous,
- TN = Number of mosquitoes correctly classified by the model as nulliparous,
- P = Total number of mosquitoes in test set that are parous, and
- N = Total number of mosquitoes in test set that are nulliparous.

. Then

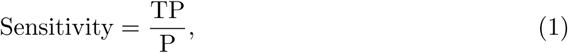

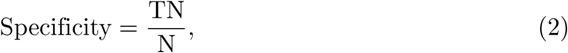

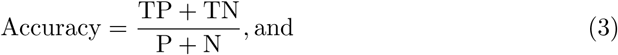

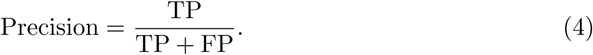

Sensitivity (also known as recall) is the percentage of correctly predicted parous mosquitoes, specificity is the percentage of correctly predicted nulliparous mosquitoes [14], and precision is the proportion of true parous mosquitoes out of all mosquitoes estimated by the model as parous [39]. We presented both sensitivity and precision because different scholars prefer one metric to another, especially for cases with imbalanced data [39]. AUC was computed from the receiver operating characteristic (ROC) curve shown in Fig 8 generated by plotting the true parous rate against the false parous rate at different threshold settings. A higher AUC is interpreted as higher predictivity performance of the model [41, 42]. The ROC curve normally presents the performance of the model at different thresholds (cut-off points), providing more information on the accuracy of the classifier [41, 42].

We hypothesized that results presented in Tables 2 and 3, and in Figs 7 and 8 were influenced by the size of a dataset used to train the model. The model that was trained on a dataset with a relatively larger number of mosquitoes, especially in the parous class, performed better than the model trained on the dataset with fewer mosquitoes.

The current standard preprocessing technique [12] leaves a mosquito spectrum with an 1851-dimensional feature space. Mathematically, binary inputs with a 1851-dimensional feature space present 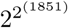 hypothesis space dimensions for the model to learn [43–45]. Successful learning of such hypothesis space dimensions requires many data points (mosquitoes in our case). Finding enough wild mosquitoes, especially parous mosquitoes, for a model to learn such a hypothesis space is expensive and time consuming. Feature reduction is an alternative to overcome this, as it reduces the hypothesis space dimension initially presented by the original data, hence lowering the number of data required to train the model efficiently. Techniques such as principal component analysis (PCA) [28, 29], partial least squares (PLS) [28, 46, 47], singular value decomposition (SVD) [31, 46, 48], and autoencoders can reduce feature space to a size that can be learned by the available data without losing important information. PCA, PLS, and SVD are commonly used when features are linearly dependent [28, 29], otherwise, an autoencoder, which can be thought as a nonlinear version of PCA, is used [30–33].

Therefore, we applied an autoencoder as illustrated in Fig 2 to reduce the spectra feature space from 1851 features to 10 features (Table 1 presents the accuracies of reconstructing original feature spaces from the encoded (reduced) feature spaces), cutting down hypothesis space dimensions from 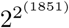 to 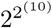, and re-trained ANN models (Figs 3 and 5).

As presented in Tables 4 - 5 and in Figs 9 - 10, the accuracy of the model improved from an average of 72% to 93%, suggesting an ANN model trained on autoencoded NIR spectra as an appropriate tool to estimate the parity status of wild mosquitoes.

**Table 4.** Performance of an ANN model trained on 75% of the encoded mosquito spectra (10 features) and tested on the remaining 25% of the encoded mosquito spectra. Minepa-ARA (Nulliparous = 656, Parous = 271), Muleba-GA (Nulliparous = 119, Parous = 21) Burkina-GA (Nulliparous = 80, Parous = 78)

|  | Minepa-ARA<br>(N=927) | Muleba-GA<br>(N=140) | Burkina-GA<br>(N=158) | Muleba-Burkina-GA<br>(N=298) |
| --- | --- | --- | --- | --- |
| Accuracy (%) | 97.1 ± 2.2 | 89.8 ± 1.7 | 93.3 ± 1.2 | 92.7 ± 1.8 |
| Sensitivity (%) | 94.9 ± 1.6 | 70.1 ± 2.3 | 91.7 ± 1.9 | 88.2 ± 2.9 |
| Specificity (%) | 98.6 ± 1.3 | 96.9 ± 1.2 | 96.4 ± 1.6 | 94.7 ± 2.1 |
| Precision (%) | 93.7 ± 2.4 | 62.5 ± 3.2 | 91.3 ± 1.4 | 93.1 ± 2.5 |
| AUC (%) | 96.7 | 91.5 | 93.1 | 94.9 |

**Figure 9.**
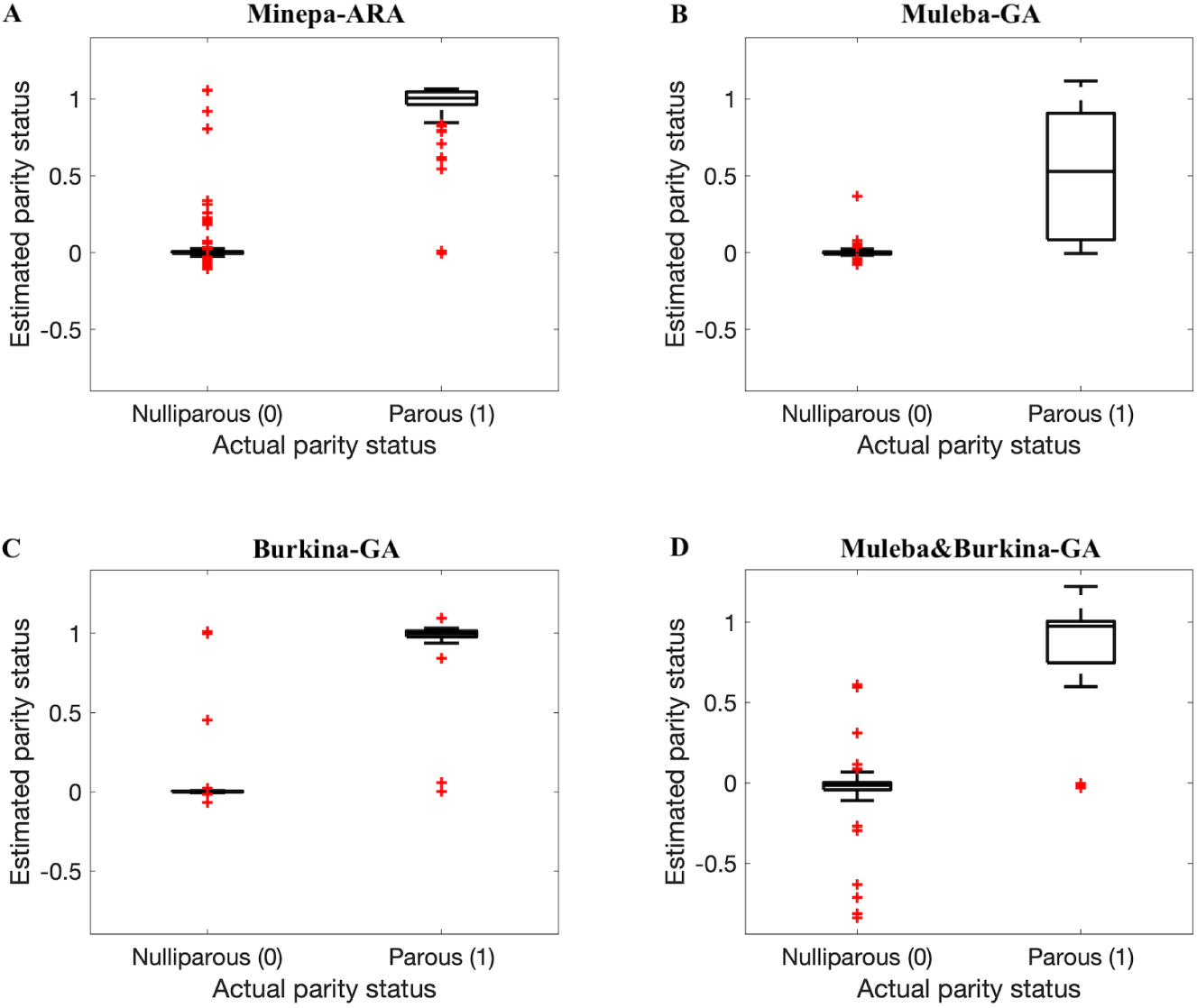
Box plots showing results when ANN models trained on 75% of encoded spectra in datasets were tested on the remaining encoded spectra (25%). A, B, C, and D represent results for Encoded-Minepa-ARA, Encoded-Muleba-GA, Encoded-Burkina-GA, and Encoded-Muleba & Burkina-GA (mosquitoes in Encoded-Muleba-GA and Encoded-Burkina-GA datasets combined) datasets, respectively

**Figure 10.**
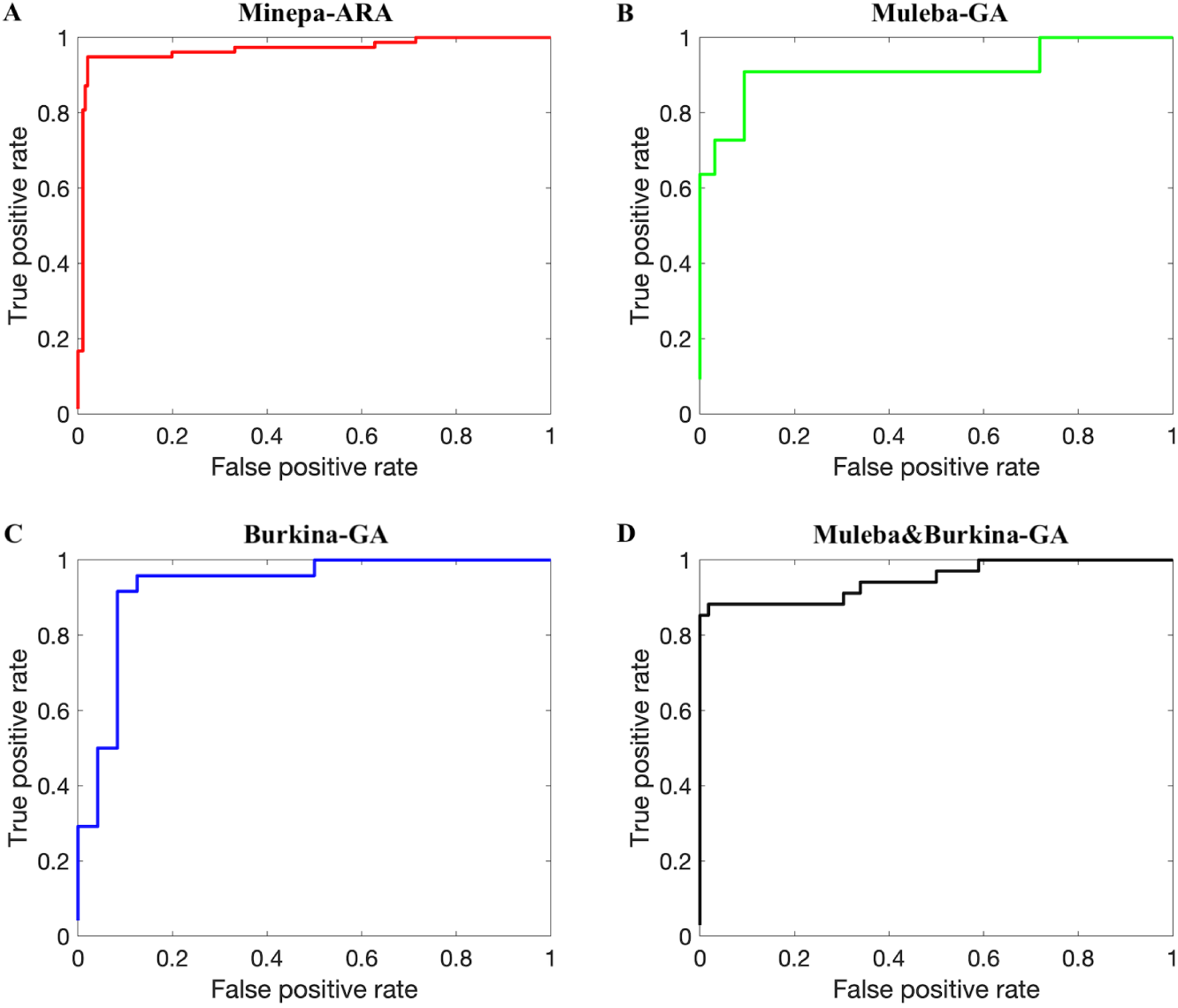
ROC curves (AUCs presented in the last row of Table 2) showing results when ANN models trained on 75% of encoded spectra were tested on the remaining encoded spectra (25%). A, B, C, and D represent results for Encoded-Minepa-ARA, Encoded-Muleba-GA, Encoded-Burkina-GA and Encoded-Muleba & Burkina-GA (mosquitoes in Encoded-Muleba-GA and Encoded-Burkina-GA datasets combined) datasets, respectively

We further applied a model trained on the Muleba-GA dataset to estimate the parity status of mosquitoes in the Burkina-GA dataset and a model trained on the Burkina-GA dataset to estimate the parity status of mosquitoes in the Muleba-GA dataset. Here we wanted to test how the model performs on mosquitoes from different cohorts. As presented in Table 6, the model performed with accuracies of 68.6% and 88.3%, respectively, showing a model trained on the Burkina-GA dataset extrapolates well to mosquitoes from a different cohort than a model trained on the Muleba-GA dataset.

**Table 5.** Confusion matrices showing accuracies of the models in absolute values when the models were trained on spectra after feature reduction by autoencoder. A) Minepa-ARA, B) Muleba-GA, C) Burkina-GA and D) Muleba-Burkina-GA. Results from the last Monte-Carlo cross validation.

|  |  | Actual Parity |  |  |
| --- | --- | --- | --- | --- |
|  | Estimates | Nulliparous | Parous | Total |
| **A** | Nulliparous | 192 | 7 | 199 |
|  | Parous | 4 | 71 | 95 |
|  | **Total** | 196 | 78 | 274 |
| **B** | Nulliparous | 33 | 2 | 35 |
|  | Parous | 3 | 5 | 8 |
|  | **Total** | 36 | 7 | 43 |
| **C** | Nulliparous | 22 | 3 | 25 |
|  | Parous | 2 | 21 | 23 |
|  | **Total** | 24 | 24 | 48 |
| **D** | Nulliparous | 58 | 4 | 62 |
|  | Parous | 2 | 27 | 29 |
|  | **Total** | 60 | 31 | 91 |

**Table 6.** Independent testing of ANN models trained on Muleba-GA and Burkina-GA encoded datasets.

|  | ANN model trained on<br>Encoded-Muleba-GA, tested on<br>Encoded-Burkina-GA | ANN model trained on<br>Encoded-Burkina-GA, tested on<br>Encoded-Muleba-GA |
| --- | --- | --- |
| Accuracy (%) | 68.6 | 88.3 |
| Sensitivity (%) | 26.5 | 86.1 |
| Specificity (%) | 94.4 | 92.2 |

A possible explanation of the results shown in Table 6 could be that, unlike for the Burkina-GA dataset, the number of parous mosquitoes (N = 21) in the Muleba-GA dataset was not representative enough for a model to learn important characteristics that extrapolate to mosquitoes in a cohort other than the one used to train the model. Although the Muleba-GA model had the poor sensitivity as presented in Table 6, the Burkina-GA model results still suggest that the ANN model trained on acceptable number of both encoded parous and nulliparous can be applied to estimate parity status of mosquitoes from different cohorts other than the one used to train the model.

## Conclusion

These results strongly suggest applying autoencoders and artificial neural networks to NIRS spectra as an appropriate complementary method to ovary dissections to estimate the parity status of wild mosquitoes. The high-throughput nature of near-infrared spectroscopy provides a statistically acceptable sample size to draw conclusions on parity status of a particular wild mosquito population. Before this method can be used as a stand-alone method to estimate parity status of wild mosquitoes, we suggest repeating of the analysis on different datasets with much larger mosquito sample sizes to test the reproducibility of the results. Hence, with the results presented in this manuscript, we recommend complementing ovary dissection with ANN models trained on NIRS spectra with their feature reduced by an autoencoder to estimate the parity status of a wild mosquito population.

## Acknowledgements

We extend our gratitude to Benjamin Krajacich for allowing us to use his already-published datasets (Burkina-GA dataset) in our analyses; the USDA, Agricultural Research Service, Center for Grain and Animal Health Research, USA for loaning us the near-infrared spectrometer used to scan the mosquitoes; and Gustav Mkandawile who worked tirelessly to make sure we obtained mosquitoes in the Minepa-ARA and Muleba-GA datasets.

Mention of trade names or commercial products in this publication is solely for the purpose of providing specific information and does not imply recommendation or endorsement by the U.S. Department of Agriculture. USDA is an equal opportunity provider and employer.

## References

1. S. Coleman, S. K. Dadzie, A. Seyoum, Y. Yihdego, P. Mumba, D. Dengela, P. Ricks, K. George, C. Fornadel, and D. Szumlas. A Reduction in Malaria Transmission Intensity in Northern Ghana After 7 Years of Indoor Residual Spraying. Malaria Journal, 16(1):324, 2017.

2. S. M. Magesa, T. J. Wilkes, A. E. P. Mnzava, K. J. Njunwa, J. Myamba, M. D. P. Kivuyo, N. Hill, J. D. Lines, and C. F. Curtis. Trial of Pyrethroid Impregnated Bednets in an Area of Tanzania Holoendemic for Malaria Part 2. Effects on the Malaria Vector Population. Acta Tropica, 49(2):97–108, 1991.

3. V. Robert and P. Carnevale. Influence of Deltamethrin Treatment of Bed Nets on Malaria Transmission in the Kou Valley, Burkina Faso. Bulletin of the World Health Organization, 69(6):735, 1991.

4. C. Dye. The Analysis of Parasite Transmission by Bloodsucking Insects. Annual Review of Entomology, 37(1):1–19, 1992.

5. C. Garrett-Jones, N. Ferreira, A. Joaquim, and W.H.O. The Prognosis for Interruption of Malaria Transmission Through Assessment of the Mosquito’s Vectorial Capacity. Technical report, Geneva: World Health Organization, 1964.

6. J. C. Beier. Malaria Parasite Development in Mosquitoes. Annual Review of Entomology, 43(1):519–543, 1998.

7. T. Detinova. Age-grouping Methods in Diptera of Medical Importance, with Special Reference to Some Vectors of Malaria. Monogr Ser World Health Organization, 47:13–191, 1962.

8. V. P. Polovodova. Age Changes in Ovaries of *Anopheles* and Methods of Determination of Age Composition in Mosquito Populations. Med Parazit (Mosk), 10:387, 1941.

9. F. E. Dowell, A. E. M. Noutcha, and K. Michel. The Effect of Preservation Methods on Predicting Mosquito Age by Near-infrared Spectroscopy. The American Journal of Tropical Medicine and Hygiene, 85(6):1093–1096, 2011.

10. B. J. Krajacich, J. I. Meyers, H. Alout, R. K. Dabiré, F. E. Dowell, and B. D. Foy. Analysis of Near-infrared Spectra for Age-grading of Wild Populations of *Anopheles gambiae*. Parasites & Vectors, 10(1):552, 2017.

11. M. F. Maia, M. Kapulu, M. Muthui, M. G. Wagah, H. M. Ferguson, F. E. Dowell, F. Baldini, and L. Cartwright. Detection of *Plasmodium falciparum* Infected *Anopheles gambiae* Using Near-infrared Spectroscopy. Malaria Journal, 18(1):85, 2019.

12. V. S. Mayagaya, K. Michael, M. Q. Benedict, G. F. Killeen, R. A. Wirtz, H. M. Ferguson, and F. E. Dowell. Non-destructive Determination of Age and Species of *Anopheles gambiae s.l.* Using Near-infrared Spectroscopy. American Journal of Tropical Medicine and Hygiene, 81:622–630, 2009.

13. V. S. Mayagaya, A. J. Ntamatungiro, S. J. Moore, R. A. Wirtz, F. E. Dowell, and M. F. Maia. Evaluating Preservation Methods for Identifying *Anopheles gambiae s.s* and *Anopheles arabiensis* Complex Mosquitoes Species Using Near-infrared Spectroscopy. Parasites & Vectors, 8(1):60, 2015.

14. M. P. Milali, M. T. Sikulu-Lord, S. S. Kiware, F. E. Dowell, G. F. Corliss, and R. J. Povinelli. Age grading *An. gambiae* and *An. arabiensis* Using Near-infrared Spectra and Artificial Neural Networks. PloS One, 14(8):e0209451, 2019.

15. M. P. Milali, M. T. Sikulu-Lord, S. S. Kiware, F. E. Dowell, G. F. Corliss, and R. J. Povinelli. Age Grading *Anopheles gambiae* and *Anopheles arabiensis* Using Near-infrared Spectra and Artificial Neural Networks. BioRxiv, page 490326, 2018.

16. A. J. Ntamatungiro, V. S. Mayagaya, S. Rieben, S. J. Moore, F. E. Dowell, and M. F. Maia. The Influence of Physiological Status on Age Prediction of *Anopheles arabiensis* Using Near-infrared Spectroscopy. Parasites & Vectors, 6(1):298, 2013.

17. M. Sikulu, G. Killeen, L. Hugo, P. Ryan, and K. Dowell. Near-infrared Spectroscopy As a Complementary Age Grading and Species Identification Tool for African Malaria Vectors. Parasite & Vectors, 3:49, 2010.

18. M. T. Sikulu, S. Majambere, B. O. Khatib, A. S. Ali, L. E. Hugo, and F. E. Dowell. Using a Near-infrared Spectrometer to Estimate the Age of *Anopheles* Mosquitoes Exposed to Pyrethroids. PLoS One, 9(3):e90657, 2014.

19. M. T. Sikulu-Lord, M. F. Maia, M. P. Milali, M. Henry, G. Mkandawile, E. A. Kho, R. A. Wirtz, L. E. Hugo, F. E. Dowell, and G. J. Devine. *Rapid and Non-destructive Detection and Identification of Two Strains of *Wolbachia* in Aedes aegypti* by Near-infrared Spectroscopy. PLoS Neglected Tropical Diseases, 10(6):e0004759, 2016.

20. M. T. Sikulu-Lord, M. P. Milali, M. Henry, R. A. Wirtz, L. E. Hugo, F. E. Dowell, and G. J. Devine. Near-infrared Spectroscopy, A Rapid Method for Predicting the Age of Male and Female Wild-type and *Wolbachia* Infected *Aedes aegypti*. PLoS Neglected Tropical Diseases, 10(10):e0005040, 2016.

21. M. P. Milali, M. T. Sikulu-Lord, S. S. Kiware, F. E. Dowell, R. J. Povinelli, and G. F. Corliss. Do NIR Spectra Collected from Laboratory-reared Mosquitoes Differ from Those Collected from Wild Mosquitoes? PloS One, 13(5).

22. C. LeClair, J. Cronery, E. Kessy, E. Tomás, Y. Kulwa, F. W. Mosha, M. Rowland, N. Protopopoff, and J. D. Charlwood. ‘Repel All Biters’: An Enhanced Collection of Endophilic *Anopheles gambiae* and *Anopheles arabiensis* in CDC Light-traps, from the Kagera Region of Tanzania, in the Presence of a Combination Mosquito Net Impregnated with Piperonyl Butoxide and Permethrin. Malaria Journal, 16(1):336, 2017.

23. T. S. Detinova. Determination of the Physiological Age of the Females of *Anopheles* by the Changes in the Tracheal System of the Ovaries. Medical Parasitology, 14(2), 1945.

24. S. M. Paskewitz and F. H. Collins. Use of the Polymerase Chain Reaction to Identify Mosquito Species of the *Anopheles gambiae* Complex. Medical and Veterinary Entomology, 4(4):367–373, 1990.

25. Q. Xu, Y. Liang, and Y. Du. Monte Carlo Cross-validation for Selecting a Model and Estimating the Prediction Error in Multivariate Calibration. Journal of Chemometrics: A Journal of the Chemometrics Society, 18(2):112–120, 2004.

26. V. S. Mayagaya, K. Michel, M. Q. Benedict, G. F. Killeen, R. A. Wirtz, H. M. Ferguson, and F. E. Dowell. Non-destructive Determination of Age and Species of *Anopheles gambiae s.l.* Using Near-infrared Spectroscopy. The American Journal of Tropical Medicine and Hygiene, 81(4):622–630, 2009.

27. A. Storkey. When Training and Test Sets are Different: Characterizing Learning Transfer. Dataset Shift in Machine Learning, pages 3–28, 2009.

28. A. M. Mouazen, B. Kuang, J. De Baerdemaeker, and H. Ramon. Comparison Among Principal Component, Partial Least Squares and Back Propagation Neural Network Analyses for Accuracy of Measurement of Selected Soil Properties with Visible and Near-infrared Spectroscopy. Geoderma, 158(1-2):23–31, 2010.

29. J. Shlens. A Tutorial on Principal Component Analysis. ArXiv Preprint 1404.1100, 2014.

30. D. Chicco, P. Sadowski, and P. Baldi. Deep Autoencoder Neural Networks for Gene Ontology Annotation Predictions. In Proceedings of the 5th ACM Conference on Bioinformatics, Computational Biology, and Health Informatics, pages 533–540. ACM, 2014.

31. B. Hervé. Auto-association by Multilayer Perceptrons and Singular Value Decomposition. Technical report, IDIAP, 2000.

32. D. P. Kingma and M. Welling. Auto-encoding Variational Bayes. ArXiv Preprint 1312.6114, 2013.

33. Y. Liu and L. Wu. High Performance Geological Disaster Recognition Using Deep Learning. Procedia Computer Science, 139:529–536, 2018.

34. C. Liou, W. Cheng, J. Liou, and D. Liou. Autoencoder for Words. Neurocomputing, 139:84–96, 2014.

35. C. Liou, J. Huang, and W. Yang. Modeling Word Perception Using the Elman Network. Neurocomputing, 71(16-18):3150–3157, 2008.

36. B. Pierre. Autoencoders, Unsupervised Learning, and Deep Architectures. In Proceedings of ICML Workshop on Unsupervised and Transfer Learning, pages 37–49, 2012.

37. D. G. Altman and J. M. Bland. Diagnostic Tests. 1: Sensitivity and Specificity. BMJ: British Medical Journal, 308(6943):1552, 1994.

38. A. G. Lalkhen and A. McCluskey. Clinical Tests: Sensitivity and Specificity. Continuing Education in Anaesthesia Critical Care & Pain, 8(6):221–223, 2008.

39. T. Saito and M. Rehmsmeier. The Precision-recall Plot is More Informative than the ROC Plot When Evaluating Binary Classifiers on Imbalanced Datasets. PloS One, 10(3):e0118432, 2015.

40. C. J. Smith. Diagnostic Tests (1)–Sensitivity and Specificity, 2012.

41. T. Fawcett. An Introduction to ROC Analysis. Pattern Recognition Letters, 27(8):861–874, 2006.

42. D. M. Powers. Evaluation: From Precision, Recall and F-measure to ROC, Informedness, Markedness and Correlation. 2011.

43. P. Stone and M. Veloso. Layered Learning. In European Conference on Machine Learning, pages 369–381. Springer, 2000.

44. P. M. Domingos. A Few Useful Things to Know About Machine Learning. Commun. ACM, 55(10):78–87, 2012.

45. T. G. Dietterich. Machine-learning Research. AI Magazine, 18(4):97–97, 1997.

46. G.H. Golub and C. Reinsch. Singular Value Decomposition and Least Squares Solutions. In Linear Algebra, pages 134–151. Springer, 1971.

47. H. Abdi. Partial Least Square Regression (PLS Regression). Encyclopedia for Research Methods for the Social Sciences, 6(4):792–795, 2003.

48. L. De Lathauwer, B. De Moor, and J. Vandewalle. A Multilinear Singular Value Decomposition. SIAM Journal on Matrix Analysis and Applications, 21(4):1253–1278, 2000.

